# A simple and reliable method for longitudinal assessment of untethered mosquito induced flight activity

**DOI:** 10.1101/2020.03.16.989897

**Authors:** Alessandro Gaviraghi, Marcus F. Oliveira

## Abstract

*Aedes aegypti* adult females are key vectors of several arboviruses and flight activity plays a central role in mosquito biology and disease transmission. Available methods to quantify mosquito flight usually require special devices and mostly assess spontaneous locomotor activity at individual level. Here, we developed a new method to determine longitudinal untethered adult *A. aegypti* induced flight activity: the INduced FLight Activity TEst (INFLATE). This method was an adaptation of the “rapid iterative negative geotaxis” assay to assess locomotor activity in *Drosophila* and explore the spontaneous behavior of mosquito to fly upon a physical stress. Insects were placed on a plastic cage previously divided in four vertical quadrants and flight performance was carried out by tapping cages towards the laboratory bench. After one minute, the number of insects per quadrant was registered by visual inspection and categorized in five different scores. By using INFLATE, we observed that flight performance was not influenced by repeated testing, sex or 5 % ethanol intake. However, induced flight activity was strongly affected by aging, blood meal and inhibition of mitochondrial complex I. This simple and rapid method allows the longitudinal assessment of induced flight activity of multiple untethered mosquitoes and may contribute to a better understanding of *A. aegypti* dispersal biology.

## 1. Introduction

Adult females of *Aedes aegypti* are facultative blood sucking insects and key vectors of several arboviruses including dengue, zika, and yellow fever viruses. Control of vector population, mainly by elimination of breeding sites and insecticide use, still represents the main strategy to prevent the spread of arboviruses. The dispersal capacity of insect vectors is a key parameter for diffusion of vector-borne diseases (Harrington et al., 2005), which is efficiently carried out by flight activity. *Aedes aegypti* mosquitoes are winged insects in their adult stages and flight activity plays multiple biological roles, required not only to seek nutrients, but also to escape hosts, mate and lay eggs in multiple sites (Muijres et al., 2017). Despite adult *A. aegypti* naturally disperse in relatively short distances from their original site, expansion to new locations poses imminent risks for human population (Guerra et al. 2014; Verdonschot and Besse-Lototskaya 2014; Honório et al., 2003; Harrington et al., 2005; Bergero et al., 2003; Leta et al. 2018). Therefore, understanding the mechanisms that mediate flight and dispersal in *A. aegypti* may drive the search for new targets for control the population of this important insect vector.

The forces required for mosquito’s take-off are generated by the orchestrated contraction of leg and indirect flight muscles (IFM), which allow the lift of engorged females after a blood meal without being detected by their hosts (Muijres et al., 2017). Mosquitoes are equipped with a pair of long and slender wings and have unique flight features. For example, mosquitoes flap their wings at very high frequencies (>700 Hz), but at very low amplitudes (~39°), which results in their characteristic sound produced by flight (Bomphrey et al., 2017). Consequently, the aerodynamic forces to lift fully engorged female mosquitoes (that usually intake about 2-3 times their own weight on blood) (Graça-Souza et al., 2006; Roitberg et al., 2003) are produced by unique kinematic mechanisms (Bomphrey et al., 2017; Muijres et al., 2017). Remarkably, recent evidence demonstrated that mosquitoes are also dispersed by means of windborne migration covering very long distances during overnight flight (Huestis et al., 2019). Notwithstanding, to reach such long distances, one must assume that mosquitoes airborne through their own flight activity and are passively carried by the wind.

The energy demand posed by flight activity is considered one of the highest known in Nature and in insects is mediated by the intense contraction of IFM (Stevenson and Josephson, 1990; Casey et al., 1985; Voigt and Winter, 1999). The energy required to complete this task is essentially provided by mitochondrial ATP production through oxidative phosphorylation (OXPHOS). Several factors regulate OXPHOS capacity and efficiency in *A. aegypti* IFM mitochondria including nutrient availability, blood digestion, sex and energy demand (Goncalves et al., 2009; Soares et al., 2015; Gaviraghi et al., 2019a; Gaviraghi et al., 2019b). However, beyond the enormous metabolic capacity of IFM to generate such extraordinary flight features of mosquitoes, several other factors determine their dispersal capacity including aging, nutritional status, blood engorgement, location, time of day, wind strength and direction, directly regulating vector competence (Harrington et al., 2005; Dor et al., 2019; Bergero et al., 2013; Huestis et al., 2019).

Available methods to quantify insect dispersal were developed to assess their spontaneous activity in the field or in laboratory settings. The mark–release–recapture (MRR) methods are very popular and make use of specific markers, such as stains or radiolabeled compounds to identify insects upon recapture (Hagler and Jackson, 2001). However, the potential toxic effects of stains used to mark individuals, and the low recovery rates, are important limitations of this methodology. Alternative approaches at laboratory settings are also available and include the flight mill to assess flight capacity and behavior of tethered insects (Krogh and Weis-Fogh, 1952; Naranjo, 2019). However, the flight aerodynamics of tethered insects are strongly affected as the power for lifting is not required given the individuals are already suspended (Taylor et al., 2010; Ribak et al., 2017). In addition, flight mill setting is time and labor-consuming requiring the tethering and following insects during flight. Regarding mosquitoes, locomotor activity was assessed using either flight mills (Rowley et al., 1968) and infrared-based computer monitor devices, firstly developed for *Drosophila* (Rosato and Kyriacou, 2006; Lima-Camara et al., 2014). Despite the benefits of these approaches, special devices and software are always required to determine spontaneous locomotor activity of mosquitoes, mostly at individual level.

Recently, a method to assess induced flight activity in male mosquitoes was designed (Culbert et al., 2018). However, this method requires a specific in-house built flight test device, beyond requiring electrical energy to induce flight and to anesthetize insects upon tests. Considering these limitations, we developed a simple, rapid and reliable method to determine longitudinal untethered adult *A. aegypti* flight activity: the INduced FLight Activity TEst (INFLATE). This method is an adaptation of the “rapid iterative negative geotaxis” assay to assess locomotor activity in *Drosophila* (Gargano et al., 2005) and explore the spontaneous behavior of mosquito to engage flight upon a physical stress (Rowley and Graham, 1968). By using this simple, rapid and reliable procedure, one can assess the induced flight activity of multiple untethered mosquitoes without the availability of electricity that can be applied anywhere, directly contributing to a better understanding of *A. aegypti* dispersal biology, and fostering the development of rational insect vector control strategies.

## 2. Material and methods

### 2.1. Ethics statement

All animal care and experimental protocols followed the guidelines of the Committee for Evaluation of Animal Use for Research of the Federal University of Rio de Janeiro (CEUA-UFRJ). The protocol was approved under the registry CEUA-UFRJ 155/13. Dedicated technicians at the Institute of Medical Biochemistry (UFRJ) animal facility performed all aspects related to rabbit husbandry under strict guidelines to ensure careful and consistent handling of the animals.

### 2.2. Insects

*Aedes aegypti* (Red eyes strain) were maintained at 28°C, 70–80% relative humidity with a photoperiod of 12 h light/dark (L:D, 12:12h). Larvae were reared on a diet consisting of commercial dog chow. Usually about 200 insects were placed in 5 L plastic cages in a 1:1 sex-ratio and allowed to feed *ad libitum* on cotton pads soaked with 10 % (w/v) sucrose solution. Since *A. aegypti* are mostly active during daytime (Gentile et al., 2009), we performed all experiments between 10 and 11 AM. Experiments were carried out using insects with either five to seven days (young group) or thirty-eight to forty-two days (old group) after the emergence. Females were fed *ad libitum* with cotton pads soaked in a 10 % sucrose solution or allowed to feed on rabbit blood. The effect of blood feeding on flight activity was assessed in five to seven days old mosquitoes fed with 10 % sucrose (control) or 24 h after a blood meal. The effect of ethanol intake on flight performance was determined by offering cotton pads soaked with either 10 % sucrose solution only (control) or supplemented with 5 % ethanol on seven days old insects. To evaluate the effect of mitochondrial complex I activity on flight ability, insects were fed for seven days on a cotton pad soaked with a 10 % sucrose solution with 5 % ethanol (control) or 250 μM rotenone dissolved in ethanol. INFLATE was performed on the seventh day after rotenone treatment. In experiments involving the use of ethanol and rotenone, the cotton pads for insect feeding were changed daily, while in the other conditions, they were changed every two days.

### 2.3. INduced FLight Activity TEst (INFLATE)

To evaluate the induced flight activity in mosquitoes, we designed the INduced FLight Activity TEst (INFLATE), which was based on the RING test used for *Drosophila* (Gargano et al., 2005; Tinkerhess et al., 2012). This method is based on the natural behavior of mosquitoes to fly upwards. An outline of the INFLATE method proposed here is schematically depicted in Figure 1. For this purpose, a plastic cage of 24 cm x 22 cm x 22 cm, was divided in four quadrants of 6 cm that have a score from 1 to 4 from the bottom to the top of the cage. At the bottom of the cage was assigned a score of zero. A group of ten mosquitoes was transferred to this cage and were allowed to acclimatize for ten minutes at room temperature. The cage was then hand-lifted to 20 cm height over a solid laboratory bench and gently tapped vertically onto its surface twice to drag down all mosquitoes to the cage bottom. The cages were then left stable onto the bench for one minute and during this time insects engaged flight activity. The number of mosquitoes present in each cage quadrant was then visually registered. To define the Induced flight activity test (INFLATE) index, the number of mosquitoes in each quadrant was multiplied by the respective score assigned to each quadrant (0, from low active to 4, from high active mosquitoes). Subsequently, the values obtained were summed and then divided by 10 (the number of mosquitoes). We then repeated this measurement for additional 19 times, allowing resting periods of two minutes between each trial. The final INFLATE index of a single experiment was calculated as the median of twenty consecutive trials, i.e. each group of 10 mosquitoes was tested twenty times to produce n = 1 of final INFLATE index.

**Figure 1.**
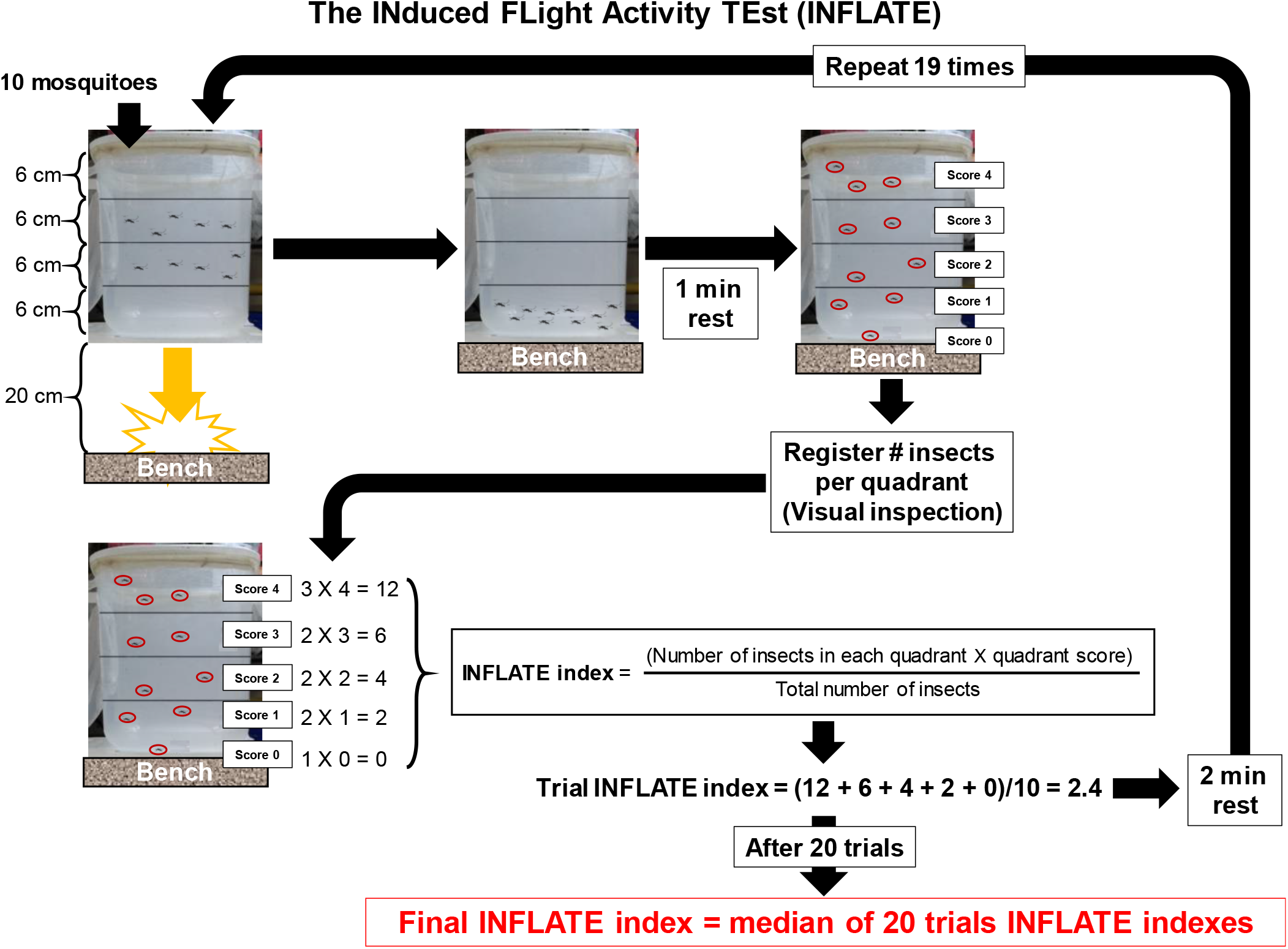
The INFLATE assay. See details in the methods section.

### 2.4. Data and statistics

Data in graphs were presented as bars with mean ± standard deviation (SD) of values for each condition. D’Agostino and Pearson normality tests were done for all values to assess their Gaussian distribution. Comparisons between groups were done by Mann Whitney’s test and differences with *p*<0.05 were considered significant. Graphs were prepared using the GraphPad Prism software version 6.00 for Windows (GraphPad Software, USA).

## 3. Results and discussion

In this work, we designed a new method to determine longitudinal untethered induced flight activity of adult *A. aegypti*: the INduced FLight Activity TEst (INFLATE). In order to validate the proposed method, experiments were carried out to evaluate the effect of several physical, chemical and physiological conditions on mosquito flight activity.

We firstly assessed if repeated tests affected the INFLATE index along the measures. Figure 2A shows the INFLATE index through twenty repetitions of young adult females fed with 10 % sucrose. We observed that INFLATE indexes were quite stable over the twenty repeats, reaching values ~ 3.5 using ten groups of ten female mosquitoes (5-9 days post-emergence) each. These experiments were performed on different days and with mosquitoes from different cohorts. Figures 2B and C show the typical distribution of mosquitoes among the quadrants of the graduated cage along the twenty repetitions of INFLATE when the experiment is conducted using young (5-7 days post-emergence, Figure 2B) and old (40-42 days post-emergence, Figure 2C) mosquitoes. We clearly observe that insect distribution among the cage quadrants in the two age classes is remarkably distinct. Indeed, quadrants 3 and 4, which indicate the highest flight capacities, are by far the most represented in young insects (Figure 2B) reaching together the 75 – 100% of insects). We also observed that no insect was found on quadrant 0 (unable to fly) and thus remained at bottom of the cage. On the other hand, quadrants 3 and 4 are less represented when old insects were challenged to fly by INFLATE (Figure 2C). In the old population, we could observe some insects that remained at bottom of the cage even after twenty repetitions of INFLATE (Figure 2C). The effect of aging on mosquito INFLATE performance will be later addressed in this work (Figure 5). Therefore, this initial set of data shows that assessment of flight activity by INFLATE is highly reproducible and not affected by repeated measures (up to twenty repetitions).

**Figure 2.**
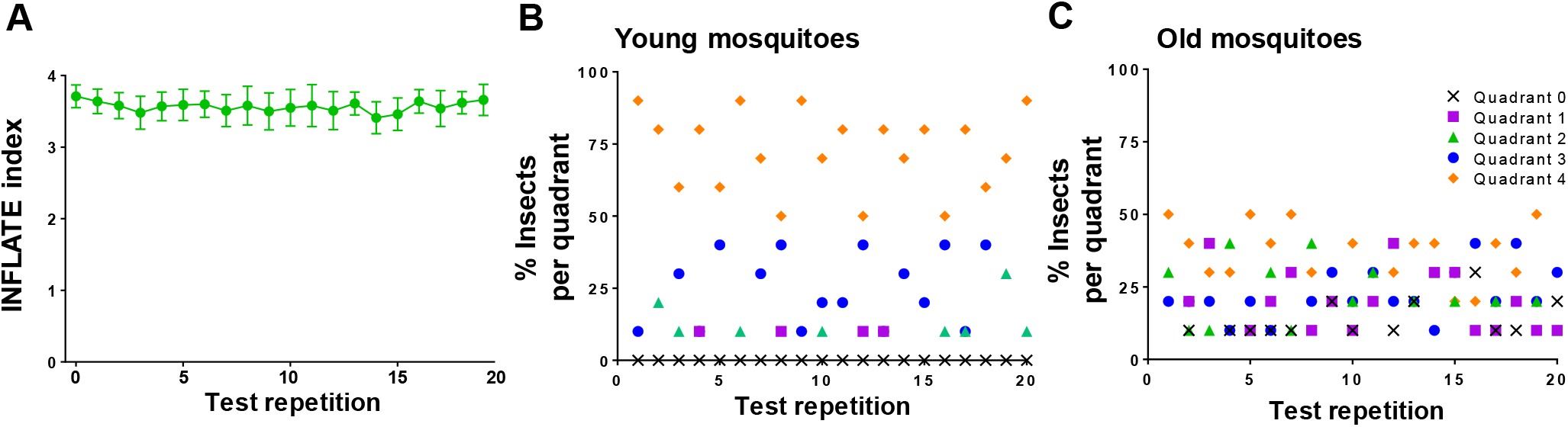
INFLATE index output is not affected by repeated measures. A) INFLATE values obtained through twenty repetitions of ten different experiments (n=10) indicate that the test shows an excellent reproducibility. B) and C) Representative scattering of insect distribution among cage quadrants from young (B) and old (C) sucrose-fed female mosquitoes. Data are expressed as mean ± standard error of the mean (SEM) of ten different experiments.

We next sought to validate INFLATE by assessing the effects of distinct physiological, chemical and physical challenges to discriminate differences in mosquito flight capacity. We thus assessed whether sex, aging, blood or ethanol intake, and inhibition of mitochondrial complex I would have any effect on INFLATE index. Figure 3 shows that INFLATE indexes obtained for young (5-7 days post-emergence) male and female mosquitoes fed exclusively with sucrose were close to 3.5, indicating very high flight capacity. Also, no significant differences were observed between the INFLATE index of males and females, indicating that induced flight activity, by using this method, is sex-independent.

**Figure 3.**
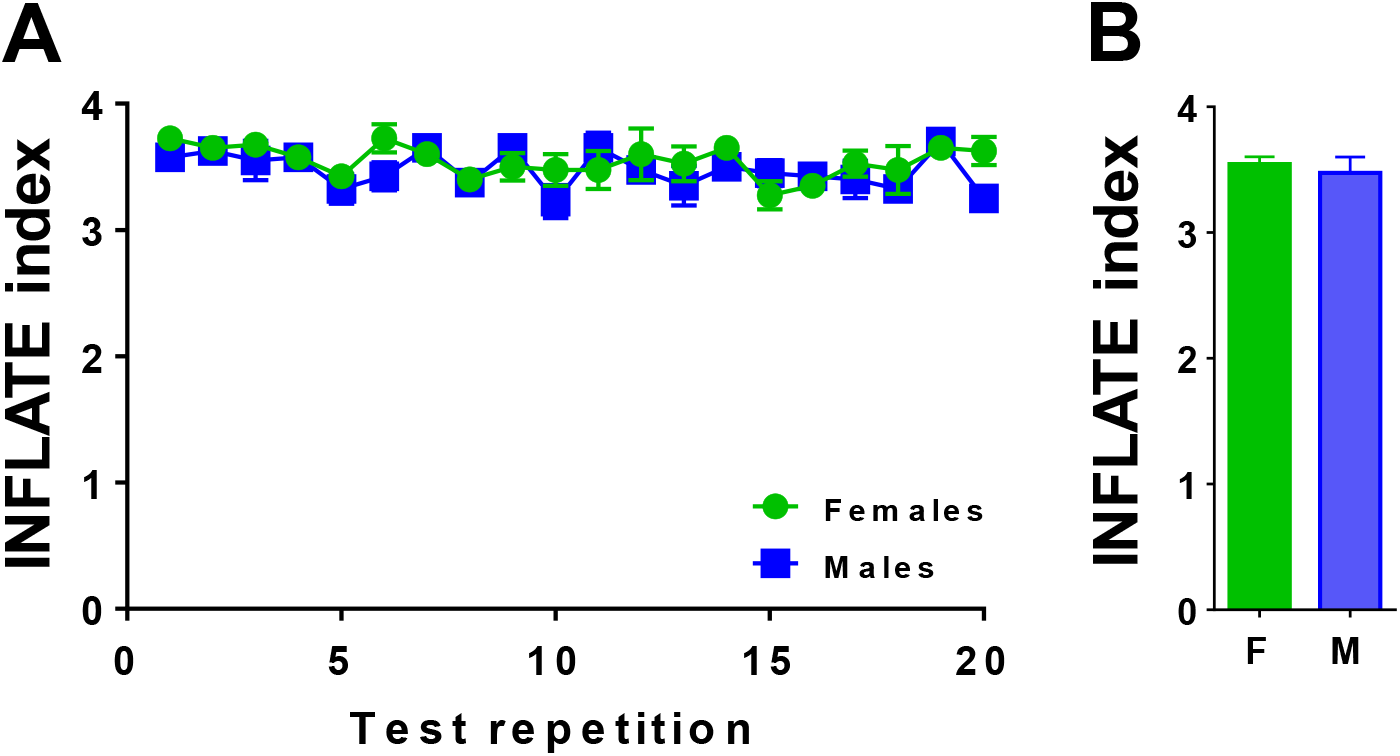
INFLATE index output is sex-independent. A) INFLATE values obtained in the twenty repetitions of four different experiments (n=4) indicate that there are no differences on induced flight activity between young sucrose-fed female and male mosquitoes. B) Compilation of INFLATE index values obtained from four different experiments (n=4). Data are expressed as mean ± standard error of the mean (SEM) of four different experiments.

This result agrees with the studies based on release-recapture experiments that report similar dispersal distance between males and females of *A. aegypti* mosquitoes (Harrington et al., 2005; Marcantonio et al., 2019). In addition, given the importance of flight in mating behavior, we anticipated that there will be no differences in flight capacity between young adult male and female mosquitoes.

In the next set of experiments, sought to test the INFLATE method in conditions where differences in flight capacity were expected. Given the huge blood engorgement capacity of adult females, we firstly tested INFLATE by analyzing the differences in flight activity between young sugar and blood-fed females using different cohorts of mosquitoes. As expected, the flight performance was remarkably affected in insects that had a blood meal 24 h before the test, reducing the INFLATE index from ~ 3.6 in sucrose-fed insect to ~ 2.4 in blood-fed ones (Figures 4A and 4B), a statistically significant decrease in flight capacity of ~ 33 %. Indeed, this result agrees with previous observations from our laboratory which demonstrated that blood feeding transiently affected flight muscle mitochondrial oxygen consumption during blood digestion (Goncalves et al., 2009). Also, after a blood-feeding, the female mosquitoes become almost totally inactive for about 48 hours (Jones, 1981; Jones and Gubbins, 1978; Lima-Camara et al., 2014). This is probably due to an inhibition in the activity of host-seeking induced by the development of the ovaries and by a mechanical stimulus induced by the abdominal distention after blood feeding. Another aspect to be taken into consideration is the enormous weight gain due to blood feeding, that usually doubles the female weight. The observed trends of INFLATE indexes along the twenty repetitions were quite constant in both groups (Figure 4A), indicating that even with a high payload of blood within their midguts female insects are still able to sustain some stable capacity to fly when challenged. Furthermore, several experiments with *A. aegypti* and other hematophagous vectors, shown that insemination and blood feeding can influence the expression of specific genes involved in circadian clocks and, consequently, on their daily activity (Meireles-Filho et al., 2006; Klowden and Lea, 1979).

**Figure 4.**
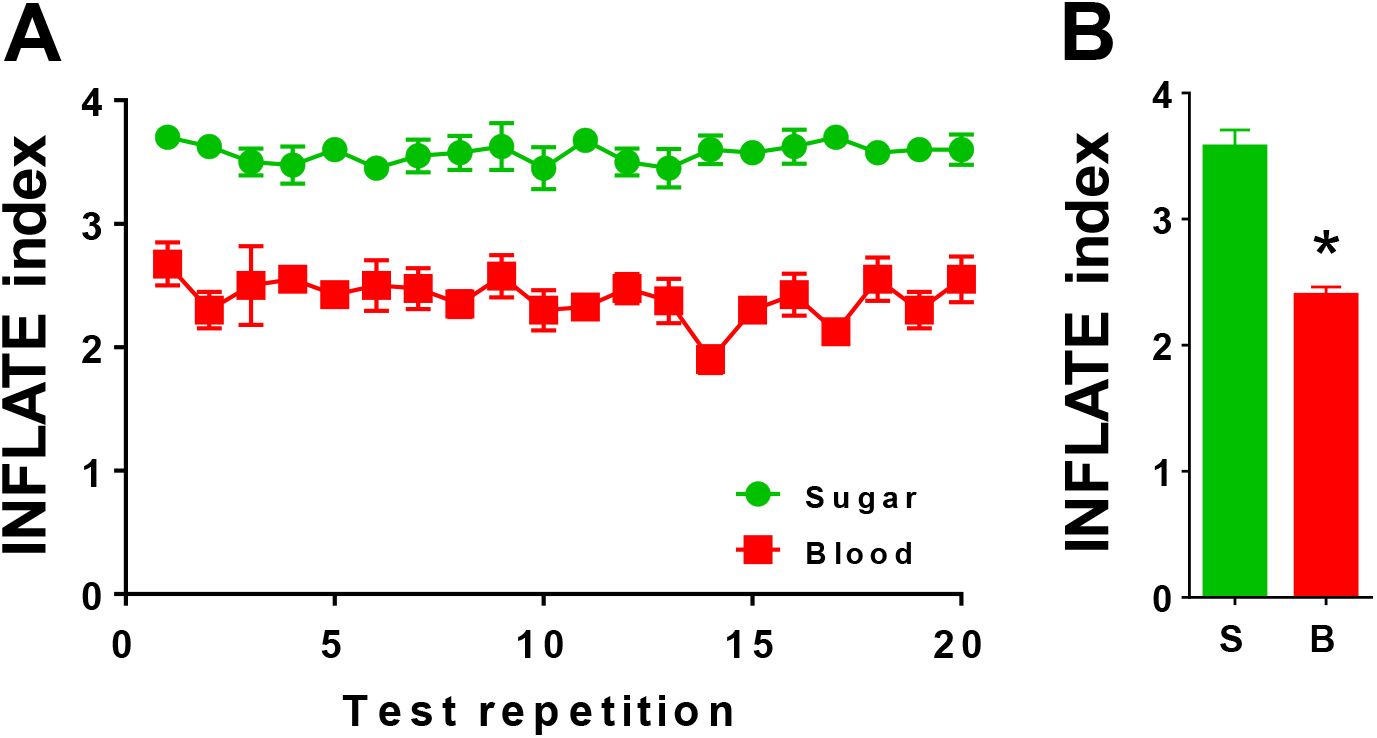
Blood-feeding reduces INFLATE index. A) INFLATE values obtained in the twenty repetitions of four different experiments (n=4) indicate that blood intake significantly reduced INFLATE index performance of young females. INFLATE was performed 24 h upon blood meal. B) Compilation of INFLATE index values obtained from four different experiments (n=4). Data are expressed as mean ± standard error of the mean (SEM) of four different experiments. Comparisons between groups were done by Mann Whitney’s test with * p < 0.05 relative to sucrose group.

We then used the INFLATE method to investigate the effects of aging on mosquito induced flight ability. Figure 5A shows that INFLATE index was remarkably low in old female insects fed exclusively with sucrose, compared to young ones. The flight performance was remarkably affected in old insects, with INFLATE indexes from ~ 3.6 in young sucrose-fed females to ~ 2.5 in old ones (Figures 5A and 5B), a statistically significant decrease in flight capacity of ~ 30 %. Interestingly, the observed reductions in old insects are quite close to those obtained upon blood engorgement in young females (Figure 4). As we previously observed (Figures 2, 3 and 4), INFLATE indexes along the twenty repetitions were quite constant regardless the aging group (Figure 5A), indicating that even with the known deterioration of muscle structure promoted by the aging process, old females are still able to sustain some stable capacity to fly when challenged (Sohal, 1976; Rockstein and Brandt, 1963; Sacktor and Shimada, 1972). These results agree with the known consequences of aging on flight capacity of different insect species (Williams et a., 1943, Lane et al., 2014; Margotta et al., 2018; Dingle, 1965; Wigglesworth, 1949; Williams et al., 1943; Rockstein and Bhatnagar, 1966; Levenbook and Williams, 1956; Tribe, 1966; Rowley and Graham, 1968; Nayar and Sauerman, 1973).

**Figure 5.**
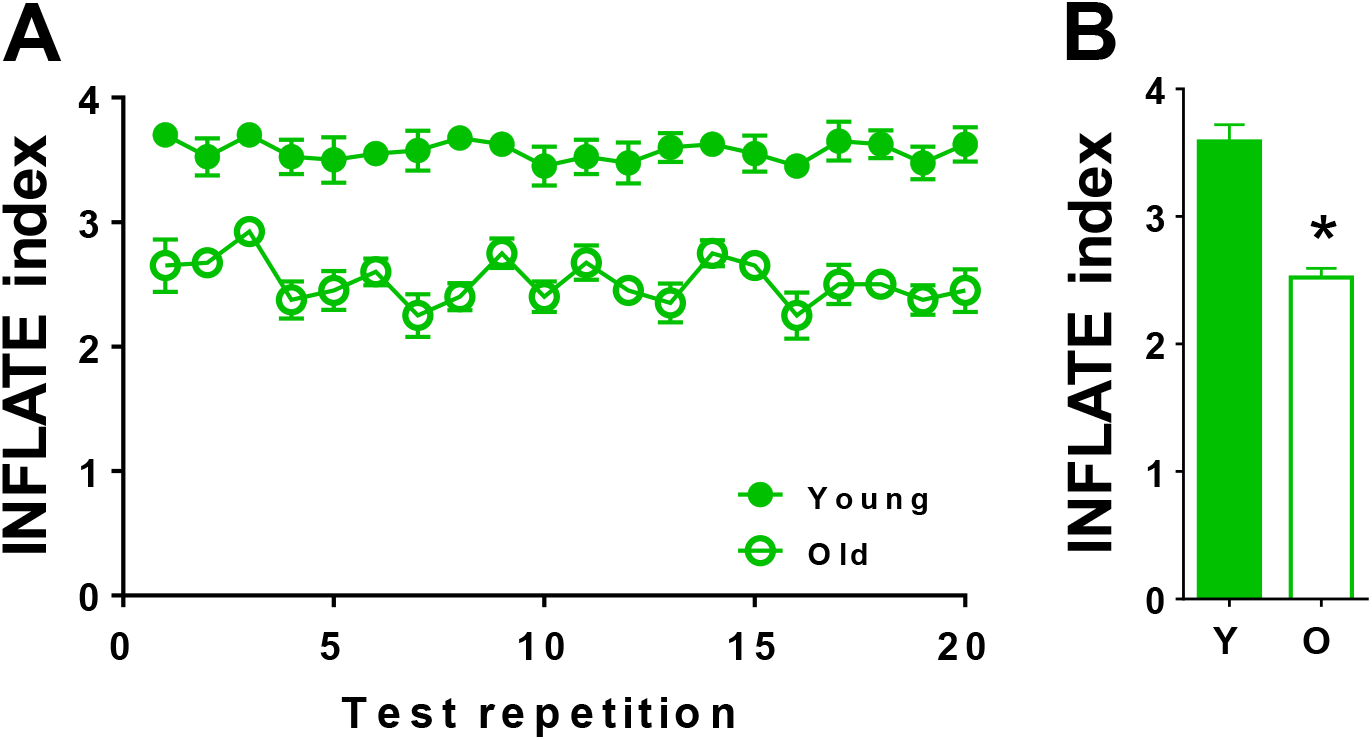
Aging affects INFLATE index. A) INFLATE values obtained in the twenty repetitions of four different experiments (n=4) indicate that aging significantly reduced INFLATE index performance of females fed exclusively with sucrose. Young insects were 5-7 days and Old insects were 38-42 days upon emergence. B) Compilation of INFLATE index values obtained from four different experiments (n=4). Data are expressed as mean ± standard error of the mean (SEM) of four different experiments. Comparisons between groups were done by Mann Whitney’s test with * p < 0.05 relative to young group.

We next tested the effects of 5 % ethanol supplementation in the diet during seven days on flight activity of sucrose-fed young females. Figure 6 shows that 5 % ethanol intake for seven days did not affect INFLATE index. Flight performance was remarkably stable along the twenty repetitions in control and 5 % ethanol insects, both groups exhibiting INFLATE indexes close to ~ 3.5 (Figures 6A and 6B).

**Figure 6.**
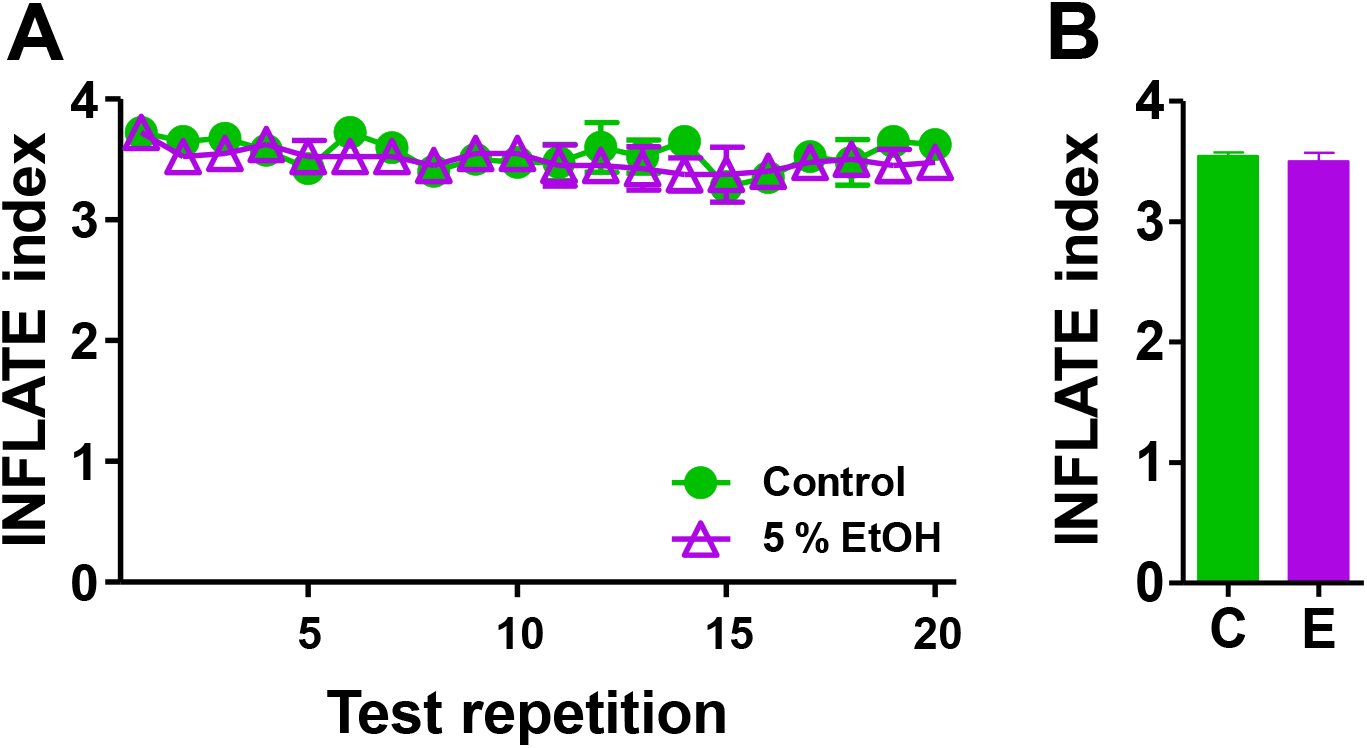
INFLATE index is not affected by dietary ethanol intake. A) INFLATE values obtained in the twenty repetitions of four different experiments (n=4) indicate that 5 % ethanol intake for seven days did not affect INFLATE index performance of females. B) Compilation of INFLATE index values obtained from four different experiments (n=4). Data are expressed as mean ± standard error of the mean (SEM) of four different experiments.

Although, to our knowledge, no direct effects of dietary ethanol supplementation were assessed on mosquito flight capacity, an interesting study revealed that ethanol and trimethylglycine (beer components) and beer itself, exhibited protective effects to mosquitoes against ionizing radiation (Rodriguez et al., 2013). Therefore, the absence of negative effects of ethanol intake on INFLATE performance (Figure 6), on survival, and in protecting against ionizing radiation (Rodriguez et al., 2013) suggest the existence of ethanol metabolizing systems in mosquitoes for proper detoxification.

Mosquito dispersal through the flight activity is a key parameter that determines vector competence, and flight muscle activity is a very high energy-demanding process. To meet this energy requirement, ATP production in flight muscles is essentially mediated by oxidative phosphorylation (OXPHOS), a process that take place in mitochondria. Multiple protein complexes located at inner mitochondrial membrane mediate a sequence of electron transfer reactions from NADH (or FADH2) to molecular oxygen. During electron transfer, part of the energy is conserved as a proton gradient across the inner mitochondrial membrane, which is ultimately used by ATP synthase complex to produce ATP. Mitochondrial complex I (or NADH:ubiquinone oxidoreductase) transfers electrons from NADH to ubiquinone and couples the energy released during the redox reactions to proton transport to intermembrane space directly contributing to ATP production by OXPHOS (Efremov et al., 2010). Rotenone, a plant derived compound largely used in the past as an insecticide, piscicide and pesticide, is a classical inhibitor of complex I (Horgan et al., 1968). Previous works from our group have shown that complex I represents one of the main sites of electron supply for ATP production in *A. aegypti* flight muscle mitochondria (Soares et al., 2015; Gaviraghi and Oliveira, 2019; Gaviraghi et al., 2019). We tested the effect of pharmacological inhibition of mitochondrial complex I activity on INFLATE performance of sucrose-fed young females. For this purpose, *we* supplemented 250 μM rotenone on sucrose diet for mosquitoes for seven days. Figure 7 shows that rotenone intake significantly reduced INFLATE index, from ~ 3.5 in control ethanol-fed insects to ~ 2.5 on complex I-inhibited insects, a statistically significant decrease in flight capacity of ~ 28 %. Flight performance was remarkably stable in control ethanol and rotenone treated insects along the twenty repetitions (Figures 7A and 7B).

**Figure 7.**
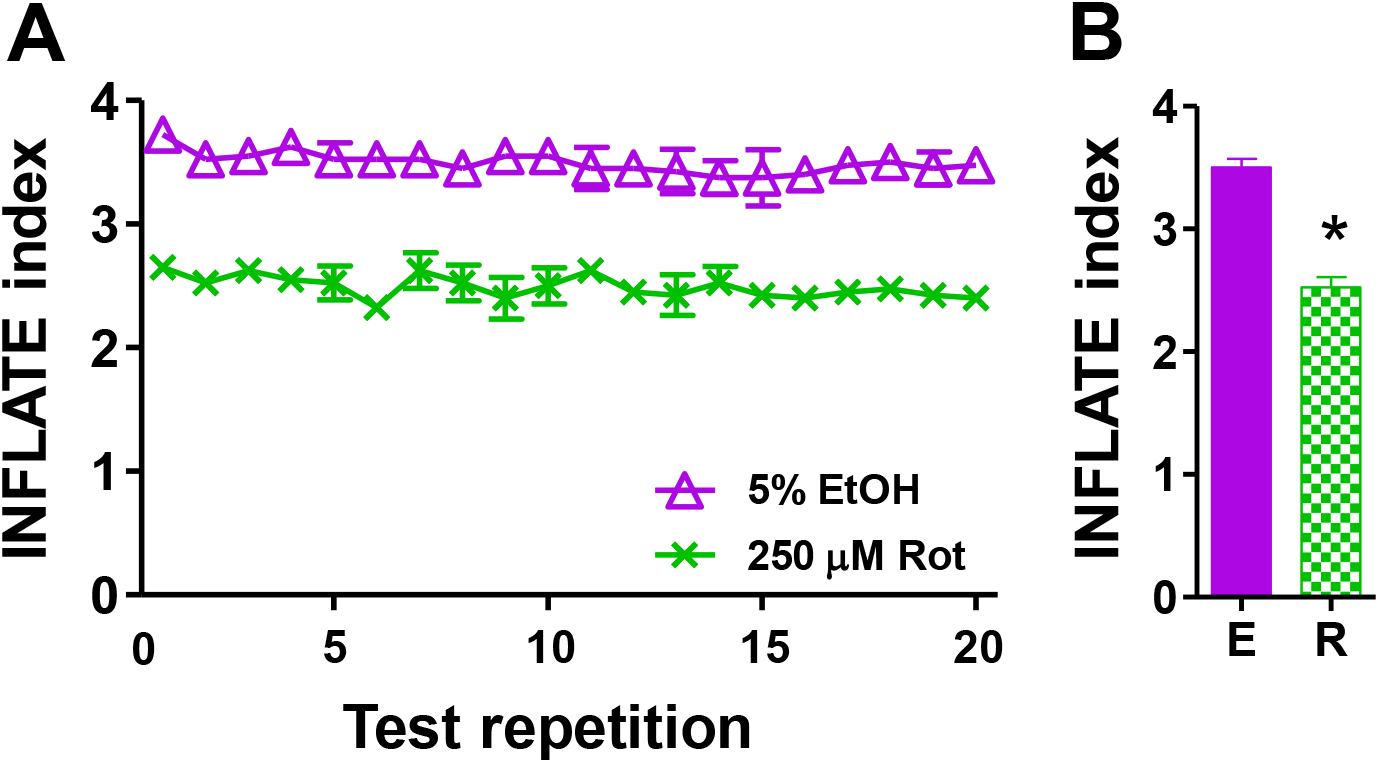
The activity of mitochondrial complex I is necessary for optimal INFLATE performance. A) INFLATE values obtained in the twenty repetitions of four different experiments (n=4) indicate that 250 μM rotenone exposure for seven days strongly affect INFLATE index performance of young females. B) Compilation of INFLATE index values obtained from four different experiments (n=4). Data are expressed as mean ± standard error of the mean (SEM) of four different experiments. Comparisons between groups were done by Mann Whitney’s test with *p<0.05 relative to EtOH group.

The main advantages that the INFLATE method offers compared to the methods currently available are: *i)* it is simple and does not require particular technical skills; *ii)* it is very cheap as only needs a graduated plastic cage; *iii)* it is fast and does not require the preparation of particular equipment and in just over an hour the results are obtained; *iv*) it does not require electricity, so it can be done in principle anywhere. Although this protocol represents a powerful tool to study mosquito flight performance, it has some limitations. Unlike the RING test used for *Drosophila* (Gargano et al., 2005; Tinkerhess et al., 2012) INFLATE has the following limitations: *i)* it is difficult to automate and depending on roughness of the plastic cage this might compromise the adequate observation of individual insects from a static point; *ii)* although a video can be taken, as is the case for free flight systems, an observer must be present to identify and record the exact location of the insect in each quadrant. Despite these technical limitations, we think that the benefits by using INFLATE outweigh the potential disadvantages described here and that this reliable, simple and cheap method represents a major step towards a better understanding of mosquito flight activity.

## 4. Conclusion

Here we report a simple, rapid and reliable method (INFLATE) to study the effect of different of several physical, chemical and physiological conditions on the induced flight activity of mosquitoes. INFLATE is a valuable tool for the systematic assessment of flight performance in several physiological and experimental conditions opening new possibilities to better understand the *A. aegypti* dispersal biology and fostering the development of rational insect vector control strategies.

## Abbreviations

RING: Rapid iterative negative geotaxis
INFLATE: Induced flight activity test
Hz: Hertz
IFM: indirect flight muscles

## 5. Acknowledgments

We would like to thank Mrs. Jaciara Miranda Freire for the excellent technical assistance on maintenance of A. aegypti colony. This study was financed in part by the Coordenação de Aperfeiçoamento de Pessoal de Nível Superior – Brasil (CAPES) – Finance Code 001, by the Conselho Nacional de Desenvolvimento Cientifico e Tecnológico (CNPq) [grant number: 404153/2016-0 MFO, and 483334/2013-8 AG], and the Fundação Carlos Chagas Filho de Amparo à Pesquisa do Estado do Rio de Janeiro (FAPERJ) [grant numbers E-26/102.333/2013, E-26/203.043/2016, and E-26/111.169/2011]. AG and MFO are fellows from CNPq [fellowship number 402409/2012-4 AG, and 303044/2017-9 MFO] and from PAPD-FAPERJ [fellowship number E-44/208702/2014]. The funders had no role in study design, data collection and analysis, decision to publish, or preparation of the manuscript.

